# Rare catechol-O-methyltransferase (COMT) missense variants are structurally unstable proteasome targets

**DOI:** 10.1101/2023.01.03.522480

**Authors:** Fia B. Larsen, Matteo Cagiada, Jonas Dideriksen, Amelie Stein, Kresten Lindorff-Larsen, Rasmus Hartmann-Petersen

## Abstract

Catechol-*O*-methyltransferase (COMT) is a key enzyme in the metabolism of catecholamines. Substrates of the enzyme include neurotransmitters such as dopamine and epinephrine, and therefore, COMT plays a central role in neurobiology. Since COMT also metabolises catecholamine drugs such as L-DOPA, variation in COMT activity could affect pharmacokinetics and drug availability. Certain COMT missense variants have been shown to display decreased enzymatic activity. Additionally, studies have shown that such missense variants may lead to loss-of-function induced by impaired structural stability, which results in activation of the protein quality control system and degradation by the ubiquitin-proteasome system. Here, we demonstrate that two rare missense variants of COMT are ubiquitylated and targeted for proteasomal degradation as a result of structural destabilisation and misfolding. This results in strongly reduced intracellular steady-state levels of the enzyme, which for the L135P variant is rescued upon binding to the COMT inhibitors entacapone and tolcapone. Our results reveal that the degradation is independent of the COMT isoform, as both soluble (S-COMT) and ER membrane-bound (MB-COMT) variants are degraded. *In silico* structural stability predictions identify regions within the protein that are critical for stability overlapping with evolutionarily conserved residues, pointing towards other variants that are likely destabilised and degraded.

## Introduction

Catechol-*O*-methyltransferase (COMT) is a key enzyme in the inactivation and metabolism of catecholamines. Substrates of COMT include those with neurotransmitter properties such as dopamine and epinephrine, and COMT has therefore been linked to multiple roles in neurobiology (Frank & Fossella, 2011; Gatt et al., 2015). The enzyme catalyses the transfer of a methyl group from *S*-adenosyl-L-methionine (SAM) to a hydroxyl group of the catechol substrate (Axelrod et al., 1958; Hanna, 1965). COMT is expressed in two major isoforms: a 221-residue soluble isoform (S-COMT) and a membrane-bound isoform (MB-COMT) containing a 50-residue N-terminal extension which forms a transmembrane region (Moo-On Huh & Friedhoff, 1979; Tenhunen et al., 1994; Ulmanen et al., 1997). The enzyme has been associated with multiple diseases such as cancer (Teng et al., 2013; Xiao et al., 2013), and cardiovascular disease (Chi Htun et al., 2011; Voutilainen et al., 2007), and COMT inhibitors are prescribed as an adjunct drug in the treatment of Parkinson’s disease (PD) (Lees, 2008; Männistö & Kaakkola, 1999).

Parkinson’s disease is amongst the most common chronic neurodegenerative disorders (Dextera & Jenner, 2013). The disease is characterised by the loss of dopaminergic neurons in the substantia nigra, consequently resulting in reduced levels of dopamine in the central nervous system (CNS) and the brain. The current symptomatic treatment involves increasing the dopamine availability. This is achieved by administering the naturally occurring dopamine precursor, L-DOPA, as dopamine itself cannot cross the blood-brain-barrier (BBB). Since the prescribed L-DOPA is a substrate of both dopamine decarboxylase (DDC) and COMT (Krauß & Bracher, 2018), the drug is sometimes administered together with both COMT (Lees, 2008) and DDC inhibitors to maximise the amount of L-DOPA transported across the BBB. However, as L-DOPA is associated with multiple side effects, it is of interest to minimise the amount of L-DOPA administered to patients (Müller, 2015), which thus indirectly may depend on the patient’s specific *COMT* alleles.

Previous studies on a range of proteins have shown that many disease-linked missense variants show loss-of-function induced by structural destabilisation of the protein (Stein et al., 2019). Many such destabilised proteins are recognised by the protein quality control (PQC) system and degraded by the ubiquitin-proteasome system (UPS) (Abildgaard et al., 2019; Ahner et al., 2007; Arlow et al., 2013; Clausen et al., 2020; Gardner et al., 2005; Kampmeyer et al., 2017b; Kriegenburg et al., 2014; Meacham et al., 2001; Nielsen et al., 2014). Previous studies of common missense variants of COMT have shown that some of these have reduced enzymatic activity and protein stability (Lachman et al., 1996; Li et al., 2005; Rutherford et al., 2008; Rutherford & Daggett, 2009; Yilmaz & Çetin, 2020). Based on these observations, we hypothesise that the observed reduction in enzymatic activity of several COMT missense variants is the result of reduced structural stability, which leads the variants to be recognised and degraded by the UPS. To address this, we studied the stability of a broader range of COMT missense variants in a cellular environment, and investigate whether loss of stability is computationally predictable.

We demonstrate that two rare missense COMT variants, as a result of structural destabilisation and misfolding, are ubiquitylated and degraded by the proteasome. The rapid proteasomal degradation results in strongly reduced steady-state levels of both the S-COMT and MB-COMT isoforms of the COMT enzyme. However, upon binding to COMT inhibitors the destabilised L135P variant is strongly stabilised and becomes soluble. Finally, we provide structure- and sequence-based saturated mutational maps and show that these *in silico* approaches help identify unstable COMT variants.

## Materials and Methods

### Buffers

Buffer A: 30 mM Tris/HCl pH 8.1, 100 mM NaCl, 5 mM EDTA, 1 mM PMSF and 1 complete mini protease. Buffer B: 30 mM Tris/HCl pH 8.1, 100 mM NaCl, 5 mM EDTA and 0.2 mM PMSF. Buffer C: 30 mM Tris/HCl pH 8.1, 100 mM NaCl, 5 mM EDTA, 2.5% Triton-X-100 and 0.2 mM PMSF. Buffer D: 20 mM Hepes pH 7.5, 0.25 M sucrose, 1 mM DTT. Buffer E: 25 mM Tris/HCl pH 8.0, 500 mM NaCl, 1 mM DTT. Buffer F: 30 mM Tris/HCl, 100 mM NaCl, 5 mM EDTA, pH 8.1. PBS: 10 mM Na_2_HPO_4_, 1.8 mM KH_2_PO_4_, 137 mM NaCl, 3 mM KCl, pH 7.4. SDS sample buffer (4x): 250 mM Tris/HCl, 8% (w/v) SDS, 50% (v/v) glycerol, 2% (v/v) β-mercaptoethanol, 0.05% bromophenol blue, pH 6.8.

### Plasmids

S- and MB-COMT cDNA were codon optimised for expression in human cells and inserted into an integrative VAMP-seq expression vector (Matreyek et al., 2018) fused to GFP (Genscript). Singlesite variants were generated by Genscript. The expression plasmid for the Bxb1 recombinase has been described before (Matreyek et al., 2018). The plasmid for expressing BFP-KDEL was from Addgene (plasmid #49150; RRID:Addgene_49150) as described before (Friedman et al., 2011). For detection of ubiquitylation, pcDNA3.1 expressing strep-Myc-tagged ubiquitin (Genscript) was used.

### Cell Culture

HEK293T landing-pad cells (Matreyek et al., 2020) were propagated in Dulbecco’s Modified Eagle High-Glucose medium (DMEM) supplemented with 10% fetal bovine serum, 5000 UI/mL penicillin, 5 mg/mL streptomycin and 2 mM glutamine at 37 □ in a humidified incubator containing 5 % CO_2_. Stable COMT transfections were performed using FugeneHD (Promega) following the manufacturer’s instructions with 3 μg COMT expression plasmid and 1 μg Bxb1 plasmid. The cells were kept for no longer than 18 passages.

Cells were treated with bortezomib (LC Laboratories) and chloroquine (Sigma) for 16 hours at concentrations of 15 μM and 20 μM, respectively. Tolcapone (Sigma) and entacapone (Sigma) were used at 10 μM for 24 hours.

### SDS-PAGE and western blotting

Unless otherwise stated samples of whole cell lysates were prepared by directly lysing in SDS sample buffer. Proteins were resolved on 7 × 8 cm 12.5 % acrylamide gels and transferred to 0.2 μm nitrocellulose membranes (Advantec, Toyo Roshi Kaisha Ltd.). Blocking of the membranes was performed with 5 % dry milk powder and 0.05 % Tween-20 in PBS. Membranes were incubated with primary antibody overnight at 4 °C followed by probing with peroxidase-conjugated secondary antibody for 1 hour. Membranes were briefly treated with ECL detection reagent (Amersham GE Healthcare) and developed using a Bio-Rad ChemiDoc MP Imaging System. The antibodies used were: anti-β-actin (diluted 1:20.000) (Sigma, A5441), anti-COMT (diluted 1:2000) (EMD Millipore Corp., AB5873-1), anti-GAPDH (diluted 1:2000) (Cell Signaling Technology, 14C10), anti-GFP (diluted 1:1000) (Chromotek, 3H9), anti-LC3a/b (diluted 1:2000) (Cell Signaling Technology, D3U4C), anti-mCherry (diluted 1:1000) (Chromotek, 6G6), anti-Myc (diluted 1:1000) (ChromoTek, 9E1), anti-Na/K-ATPase (diluted 1:2000) (EMD Millipore Corp., C464.6), anti-calreticulin (diluted 1:2000) (Invitrogen, PA5-25922), anti-mouse (diluted 1:5000) (Dako, P0260), anti-rabbit (diluted 1:5000) (Dako, P0217) and anti-rat (diluted 1:5000) (Invitrogen, 31470).

### Solubility experiments

Cells were lysed on ice in buffer A followed by sonication 3 × 20 seconds. Cell lysates were centrifuged at 15,000 g at 4 °C for 30 minutes. The supernatant and pellet fractions were separated, and the pellet was resuspended in a volume of buffer A identical to the volume of the supernatant. Then, SDS sample buffer was added and the samples analysed by SDS-PAGE and western blotting.

### Immunoprecipitation

Stable COMT cell lines were transfected with Myc-tagged ubiquitin using FugeneHD (Promega) following the manufacturer’s instructions using 4 μg plasmid. The day after transfection, the cells were treated with 10 μM bortezomib (LC Laboratories) for 16 hours. After incubation, the cells were harvested on ice in 300 μL buffer B followed by sonication 3 × 10 seconds. After sonication, 75 μL 8 % SDS was added and the cell lysates were boiled for 10 minutes and vortexed. The samples were left to rest for 5 minutes at room temperature followed by addition of 1125 μL buffer C. The samples were left on ice for 30 minutes and centrifuged at 16,000 g for one hour at 4 °C and the protein was captured with 25 μL Myc-trap (Chromotek) beads by tumbling overnight. The beads were washed by centrifugation in buffer F five times and the proteins were eluted in 40 μL SDS sample buffer followed by boiling for 2 minutes. The samples were analysed by SDS-PAGE and western blotting.

### Fluorescence microscopy

For fluorescence microscopy, cells were treated with bortezomib and tolcapone as described above and analysed by live cell imaging on a Zeiss AxioVert microscope equipped with a 10x objective and a digital camera (Carl Zeiss AxioCam ICm1). For co-localisation experiments, stable HEK293T cells expressing wild type COMT were transiently transfected with pTagBFP-C expressing the KDEL marker using FugeneHD (Promega) following the manufacturer’s instructions with 100 ng plasmid. Two days after transfection, the cells were transferred to a 96-well uClear plate (Greiner) and incubated for 24 hours. Then, the cells were imaged on a Molecular Devices ImageXpress Confocal HT.ai microscope.

### Flow cytometry

Cells were dislodged with trypsin, washed with PBS and resuspended in PBS with 2 % fetal bovine serum followed by filtering through a 35 μm nylon mesh filter. The cells were examined using a BD FACSJazz flow cytometry machine and the BD FACS Software 1.2.0.142 and analysed with FlowJo 10.1r1 software (BD Biosciences).

### MB-COMT membrane topology

Cells were harvested and lysed on ice in buffer D followed by centrifugation at 5000 g for 2 minutes. The cell pellet was resuspended in buffer D using a 27-gauge needle followed by centrifugation at 300 g for 3 minutes at 4 □ to discard unbroken cells. The supernatant was transferred to a fresh tube and centrifuged at 100,000 g for 60 minutes at 4 □. The cell pellet was resuspended in buffer E and aliquots were treated on ice with triton X-100 and/or proteinase K for 60 minutes. Finally, proteinase K was inhibited by adding 5 mM PMSF. Then, the samples were subjected to precipitation with trichloroacetic acid (TCA). Briefly, 15% TCA was added to the samples followed by centrifugation at 17,000 g for 30 minutes at 4 □. The precipitates were washed in ice cold acetone and resuspended in SDS sample buffer. The samples were analysed by SDS-PAGE and western blotting.

### Structural stability predictions

Changes in thermodynamic stability of COMT variants (ΔΔG) were predicted using Rosetta (GitHub SHA1 99d33ec59ce9fcecc5e4f3800c778a54afdf8504) with the Cartesian ddG protocol (Park et al., 2016) using the COMT crystal structure (PDB: 5LSA) as input after removing other molecules in the PDB file. The ΔΔG values obtained from Rosetta were divided by 2.9 to convert them from Rosetta energy units to a scale corresponding to kcal/mol (Park et al., 2016).

### Evolutionary conservation scores

An *in silico* evolutionary distance was calculated for all the MB-COMT variants using information from evolutionary sequence conservation. A multiple sequence alignment of 1532 MB-COMT homologs was generated through HHblits (Remmert et al., 2012) using an E-value threshold of 10^−20^.

By filtering out sequences with more than 50% gaps, this dataset was reduced to 1490 homologs. From this, evolutionary conservation scores were calculated using the Global Epistatic Model for predicting Mutational Effects (GEMME) software (Laine et al., 2019).

## Results

### *In silico* saturation mutagenesis and thermodynamic stability predictions

To explore the mutational landscape of COMT variants we first employed structure-based energy calculations to predict the effect of amino acid substitutions on the thermodynamic folding stability of the COMT enzyme. *In silico* saturation mutagenesis was performed by introducing all possible single-site amino acid substitutions into the wild type crystal structure of COMT (Ehler et al., 2014). Then, using the Rosetta energy function (Park et al., 2016), the difference (Δ) in thermodynamic folding stability (ΔG) between the variants and wild type COMT (ΔΔG) was predicted for all 19 possible variants at each position resolved in the COMT crystal structure (PDB: 5LSA). As the structure does not provide information on the N-terminal transmembrane region (in MB-COMT) and last five residues (in both S- and MB-COMT) we could not predict the effects of missense variants in these regions. In total the dataset comprises 4047 different COMT missense variants. The entire dataset is included in the supplemental material (Supplemental File 1) and presented here as a heat-map (Fig. 1A). As the ΔΔG values report on the difference in folding stability, low ΔΔG scores (white/yellow colours in Fig. 1A) indicate substitutions predicted not to cause any major disturbances to the enzyme structure or stability. Conversely, high ΔΔG scores (orange/red colours in Fig. 1A) predict a destabilisation of the COMT structure, possibly resulting in degradation by the UPS in a cellular context.

**Fig. 1.**
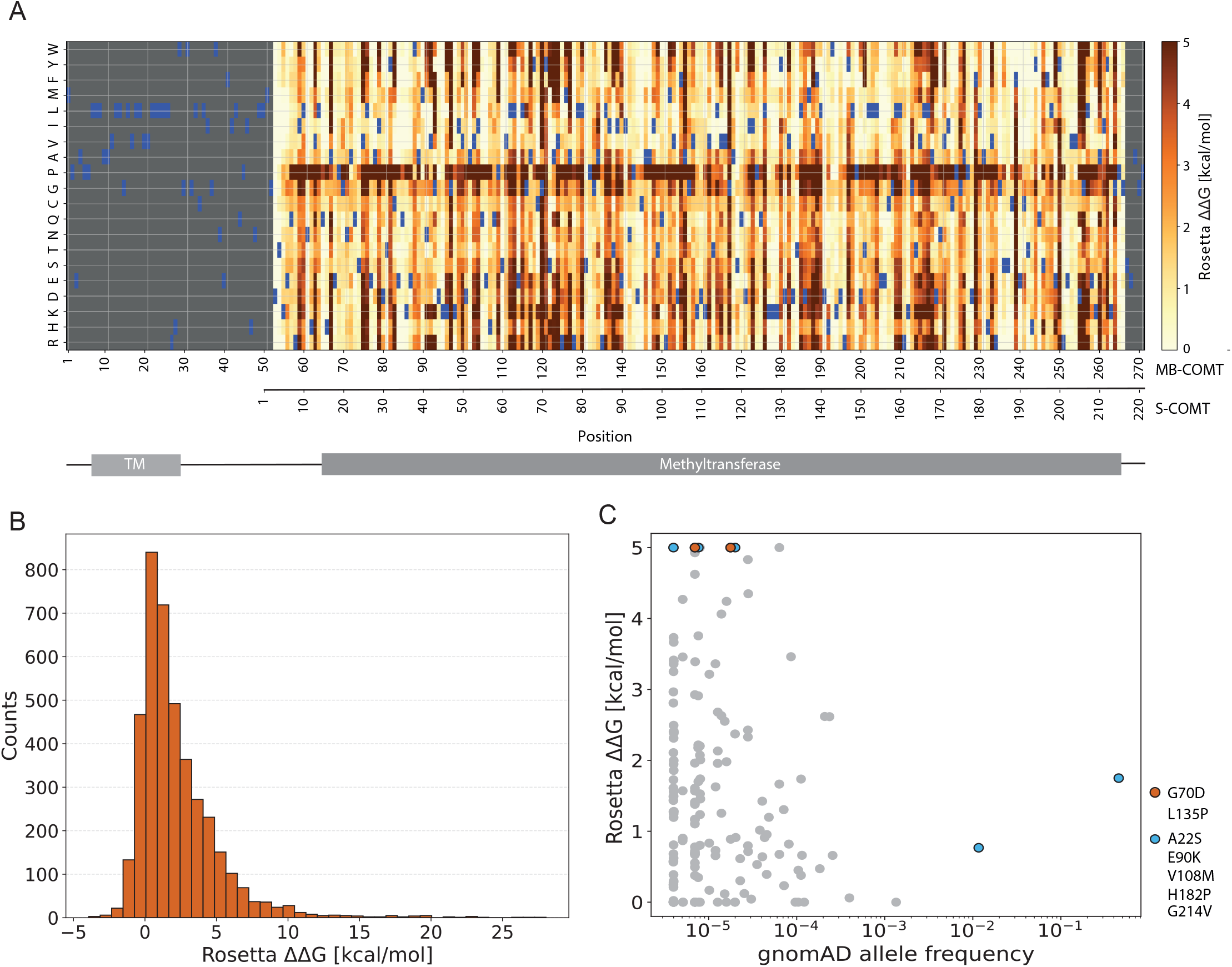
*In silico* thermodynamic stability predictions. (A) Heatmap of predicted thermodynamic stability (ΔΔG) of COMT variants. *In silico* saturation mutagenesis introducing all possible singlesite amino acid substitutions into the COMT crystal structure (PDB: 5LSA) (Ehler et al., 2014) and using Rosetta to predict the thermodynamic stability (ΔΔG) for each variant. As the 50-residue N-terminal region and the last 5 residues are not resolved in the crystal structure, we could not perform the calculations for these residues. Wild type (WT) residues are indicated in blue. The numeration of S-COMT and MB-COMT is indicated, as well as the transmembrane (TM) and methyltransferase domain in the COMT structure (Letunic et al., 2021). (B) The distribution of Rosetta ΔΔG scores of COMT variants. (C) The Rosetta ΔΔG scores of variants reported in the Genome Aggregation Database (gnomAD) (Karczewski et al., 2020). The two variants that we find to have low cellular abundance G70D/G120D (S-COMT/MB-COMT) and L135P/L185P (S-COMT/MB-COMT), are indicated in red, and the variants that have a cellular abundance at least half of the wild-type protein (A22S, E90K, V108M, H182P, G214V (S-COMT)) are indicated in blue.

As expected, and in agreement with deep mutational scans (Gray et al., 2017; Høie et al., 2022) and Rosetta-based predictions (Abildgaard et al., 2019; Cagiada et al., 2021; Nielsen et al., 2021) on a range of proteins, substitutions to proline appears detrimental at most positions (Fig. 1A). In general, many amino acid substitutions are relatively well tolerated (Fig. 1B), while some variants lead to a predicted decrease in stability (increased ΔΔG). For example, we find 41% of the variants have predicted values of ΔΔG > 2.0 kcal/mol and 15% have ΔΔG > 4.5 kcal/mol. These cut-off values have previously been shown to correlate with moderate to substantial loss of stability and function (Høie et al., 2022), and show enrichment in disease-causing missense variants (Blaabjerg et al., 2022).

As a first step to test the validity of the stability predictions, we compared the ΔΔG scores of variants reported in the Genome Aggregation Database (gnomAD) (Karczewski et al., 2020). In line with a previous analysis of a large set of proteins (Blaabjerg et al., 2022), this revealed that variants observed at a high frequency within the population all display low ΔΔG values suggesting that these common alleles are structurally stable and functional; for example the variants with allele frequencies > 0.001 all have ΔΔG < 2.0 kcal/mol (Fig. 1C). However, the rarer COMT variants displayed a wide variety of ΔΔG scores, including several with high ΔΔG values (Fig. 1C), suggesting that they potentially could be targets of the cellular PQC system.

### Unstable S-COMT variants are targeted for proteasomal degradation

To test if destabilised COMT variants were indeed targeted for PQC-linked proteasomal degradation we selected wild type S-COMT and seven missense variants with predicted ΔΔG values ranging from 0.8 to 8.9 kcal/mol (Table 1) for further experimental analysis. It should be noted that the residue numeration of S-COMT, lacking the first 50 N-terminal residues, is initialised at MB-COMT residue M51 (Fig. 1A). Our selection includes previously characterised variants such as S-COMT A22S and V108M, both associated with reduced enzymatic activity (Chi Htun et al., 2011; Rutherford & Daggett, 2009; Teng et al., 2013; Voutilainen et al., 2007; Xiao et al., 2013), as well as rarer variants reported only in gnomAD. In addition, the variants were selected to ensure that the positions of the substitutions were distributed throughout the COMT sequence and structure (Fig. 2A).

**Table 1.** Selected COMT variants.

| S-COMT<br>variant | MB-COMT<br>variant | gnomAD allele<br>frequency | S-COMT Rosetta<br>$\Delta\Delta G$ (kcal/mol) | GEMME $\Delta E$<br>score |
| --- | --- | --- | --- | --- |
| WT | WT | ~0.50 | 0 | 0 |
| A22S | A72S | 0.0136 | 0.77 | -1.9 |
| G70D | G120D | $3.19 \times 10^{-5}$ | 8.6 | -7.0 |
| E90K | E140K | $2.00 \times 10^{-5}$ | 5.4 | -5.8 |
| V108M | V158M | 0.46 | 1.8 | -1.2 |
| L135P | L185P | $7.95 \times 10^{-6}$ | 6.8 | -3.8 |
| H182P | H232P | $3.98 \times 10^{-6}$ | 6.7 | -1.4 |
| G214V | G264V | $8.02 \times 10^{-6}$ | 6.0 | -2.7 |

**Fig. 2.**
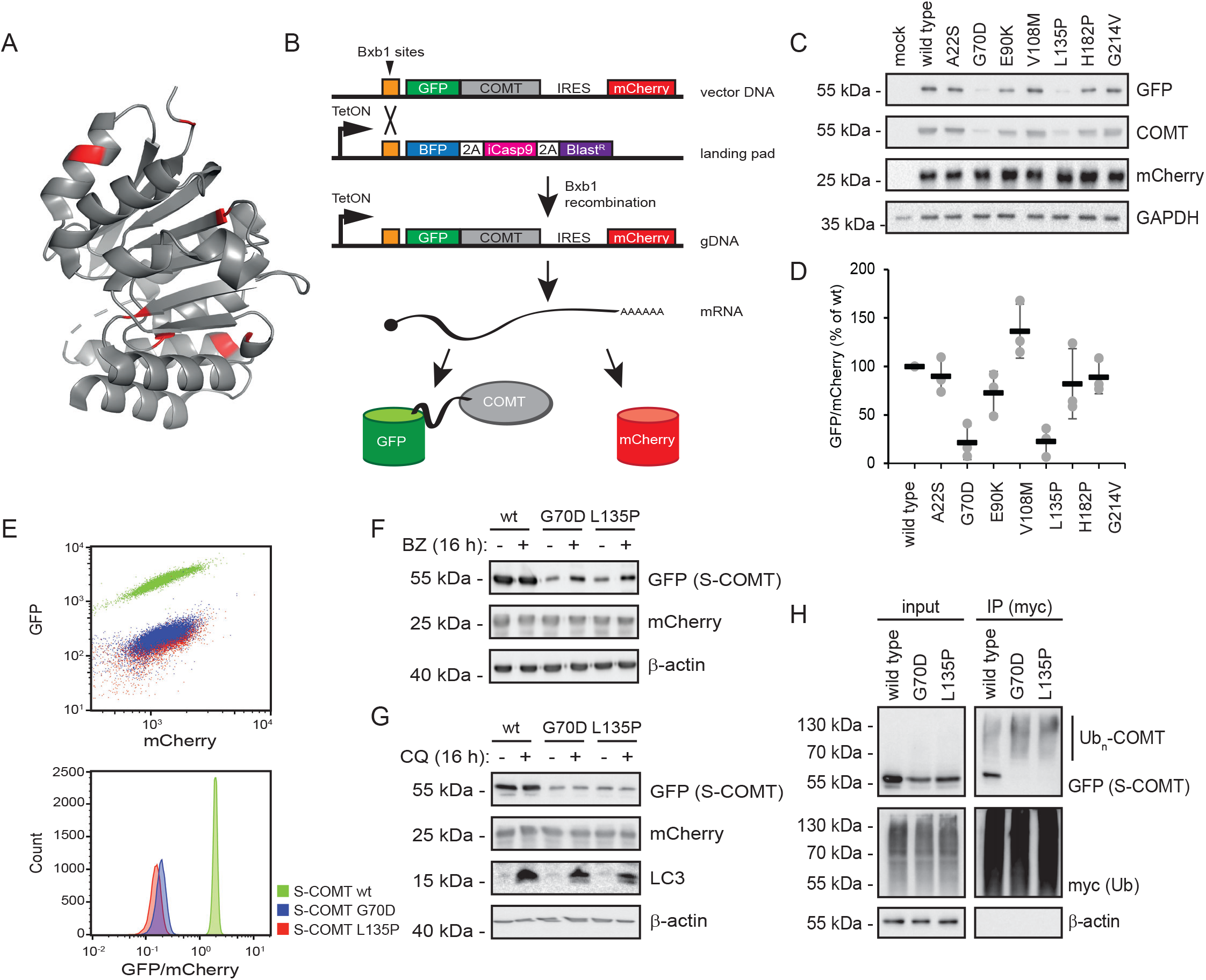
Unstable S-COMT variants are targeted for proteasomal degradation. (A) The crystal structure of S-COMT (PDB: 4PYI). The seven mutated residues are indicated in red. The figure was made in PyMOL. (B) S-COMT variants are N-terminally fused to GFP followed by an internal ribosome entry site (IRES) and the reporter gene, mCherry. The plasmid is introduced into HEK293T cells containing a landing-pad site expressing a blue fluorescent protein (BFP), two 2A-like translational stop-start sequences, inducible caspase 9 (iCasp9) and blasticidin S (Blast). Upon Bxb1 recombination, the plasmid is introduced into the landing-pad site and the *BFP-iCasp9-Blast^R^* will be displaced further downstream away from the promoter (not depicted), thus inhibiting expression. From the genomic DNA (gDNA) landing-pad site mRNA is expressed, which is translated into two individual proteins, GFP-COMT and mCherry. Expression is controlled by a tetracycline-regulated promoter (Tet-ON). (C) The steady-state protein levels of wild type (WT) S-COMT and the seven S-COMT variants in HEK293T whole cell lysates determined by SDS-PAGE and western blotting using antibodies against GFP and COMT. Antibodies against mCherry and GAPDH were included as controls for equal expression and loading, respectively. (D) Quantification of blots as shown in (A) by densitometry normalised to the mCherry signal. The WT is set to 100%. Error bars indicate the standard deviation (n=3). (E) Scatter plot (upper panel) showing the quantification of the GFP and mCherry signal of HEK293T cells expressing S-COMT WT, G70D and L135P by flow cytometry. Histogram (lower panel) showing the GFP/mCherry ratio. (FG) Protein levels in whole cell lysates of HEK293T cells expressing S-COMT WT, G70D and L135P with or without treatment with proteasomal inhibitor bortezomib (BZ) (F) or autophagy inhibitor chloroquine (CQ) (G) for 16 hours. Antibodies against β-actin were included as a loading control and antibodies against the autophagy substrate, LC3, were included as a positive control for CQ treatment. (H) Denaturing immunoprecipitation (IP) of ubiquitin (Ub) using myc-trap resin and myc-tagged ubiquitin (Ub). The precipitated material was analysed by SDS-PAGE and western blotting using antibodies against GFP (S-COMT) and myc (ubiquitin). β-actin was included as a control.

The variants were expressed from an integrative mammalian expression vector as GFP fusion proteins. To account for cell-to-cell variations in expression, mCherry was co-transcriptionally expressed with COMT and translated from an internal ribosomal entry site (IRES) (Fig. 2B). The variants were introduced into HEK293T cells containing a Bxb1 landing-pad site to ensure site-specific integration of the variants (Matreyek et al., 2018, 2020). As the expression plasmid does not contain any promoter, single copy tetracycline/doxycyclin-regulated expression is achieved from the Tet-ON promoter at the landing-pad locus (Fig. 2B).

When we compared the steady-state levels of the selected S-COMT variants by western blotting, two of the variants, G70D and L135P, displayed strongly reduced steady-state levels (Fig. 2C), while the remaining variants, including the common alleles A22S and V108M, appeared like wild type (WT) S-COMT (Fig. 2C). Quantifications revealed that the G70D and L135P variants were significantly reduced by at least 4 to 5 fold (Fig. 2D); the differences in steady-state levels were about 10 fold when we instead compared the levels by flow cytometry (Fig. 2E).

To investigate if the reduction in cellular abundance of S-COMT G70D and L135P is indeed a result of UPS-mediated degradation, we compared the levels of the variants after treatment with the proteasome inhibitor bortezomib (BZ). Indeed, this led to an increased level of both G70D and L135P, whereas wild type S-COMT was unaffected (Fig. 2F). Conversely, addition of the autophagy inhibitor, chloroquine (CQ), did not affect any of the tested variants (Fig. 2G). Additionally, denaturing immunoprecipitation experiments to probe the ubiquitylation pattern of the variants revealed that although some wild type S-COMT appears to be modified with a single ubiquitin moiety, the unstable variants appeared more heavily modified with multiple ubiquitin molecules (Fig. 2H).

Finally, when we fused the GFP-tag to the C-terminus of S-COMT, this did not affect the proteasome-dependent reduced steady-state levels (Fig. S1), showing that the destabilisation is independent of the position of the GFP fusion. Based on these experiments, we conclude that the unstable S-COMT variants are targeted for ubiquitin-dependent proteasomal degradation.

### COMT inhibitors stabilise the L135P variant

A number of previous studies have indicated that the PQC system may be over-meticulous, thus targeting variants that are only slightly destabilised and including some that might still be functional (Gardner et al., 2005; Kampmeyer et al., 2022; Kampmeyer et al., 2017a; Kriegenburg et al., 2014; Meacham et al., 2001). Thus, perhaps similar to disease-linked missense variants in the protein DHFR, which are dramatically stabilised by the DHFR inhibitor methotrexate (Kampmeyer et al., 2022), certain COMT variants could potentially be stabilised by COMT inhibitors. To test this, cells expressing wild type S-COMT or the G70D and L135P variants were treated with the well-characterised and clinically approved COMT inhibitors, entacapone (ENT) and tolcapone (TOL) (Lees, 2008). This revealed no effect on the cellular abundance of wild type or G70D S-COMT. However, for L135P we observed strongly increased levels upon treatment with either of the inhibitors, indicating that this variant is structurally stabilised when bound to the inhibitors (Fig. 3AB). Again this effect was independent on whether the GFP-tag was fused to the N-terminus (Fig. 3) or C-terminus (Fig. S1B) of S-COMT.

**Fig. 3.**
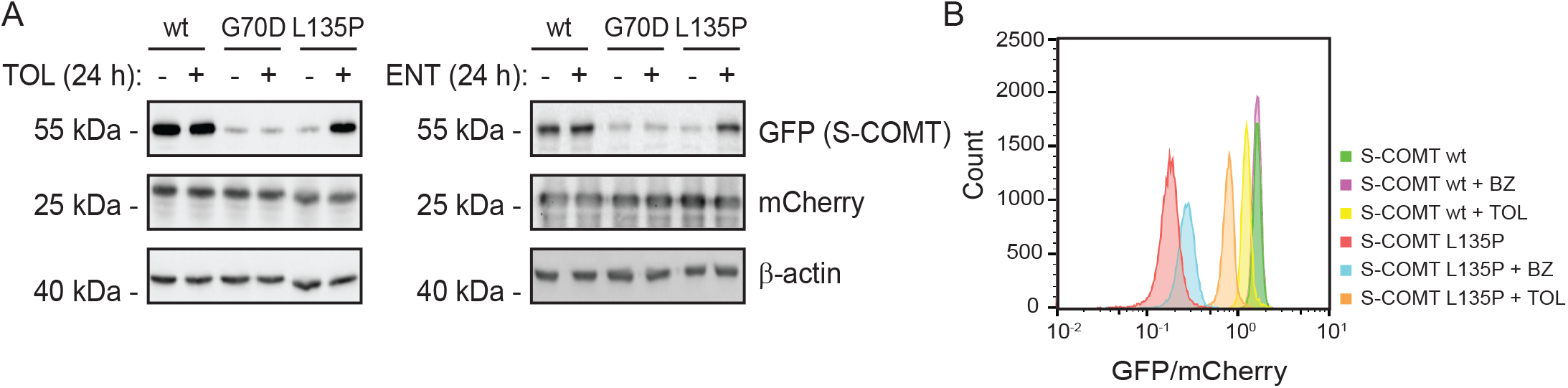
COMT inhibitors stabilise the S-COMT L135P variant. (A) The steady-state protein levels of whole cell lysates of HEK293T cells expressing wild type (WT) S-COMT, G70D and L135P with or without incubation with COMT inhibitors tolcapone (TOL) or entacapone (ENT) for 24 hours. Antibodies against mCherry and β-actin were included as controls for equal expression and loading, respectively. (B) Quantification by flow cytometry of the GFP/mCherry ratio of HEK293T cells expressing WT S-COMT and L135P after incubation with the proteasomal inhibitor bortezomib (BZ) or the COMT inhibitor tolcapone (TOL).

### Aggregation and cellular localisation of COMT variants

As destabilised proteins are prone to misfold and form insoluble aggregates, we next analysed the subcellular localisation of the variants by fluorescence microscopy. The wild type, G70D and L135P variants all localised to the cytosol (Fig. 4A). However, as expected, the cells expressing the unstable variants showed clearly reduced GFP intensities compared to wild type S-COMT (Fig. 4A). Corroborating the results above, treatment with tolcapone did not affect the G70D GFP signal, while for the L135P variant an increased signal was observed (Fig. 4A). Conversely, after adding bortezomib both G70D and L135P levels were increased (Fig. 4A), but rather than being evenly distributed in the cytosol, both variants now localised in large cytosolic inclusions (Fig. 4A). This indicates that while the tolcapone-mediated increase in the L135P steady-state level is a result of the L135P variant becoming structurally stabilised and thus averting PQC-linked degradation, the proteasome-inhibition, on the other hand, leads to an increase in non-native aggregation-prone S-COMT G70D and L135P protein species. Accordingly, when we compared the solubility of the wild type, G70D and L135P S-COMT variants by differential centrifugation and western blotting, G70D and L135P were mainly found in the insoluble (pellet) fractions (Fig. 4B). Blocking the proteasome with bortezomib did not affect the solubility of the G70D and L135P variants, while addition of tolcapone led to an increased amount of soluble L135P (Fig. 4B).

**Fig. 4.**
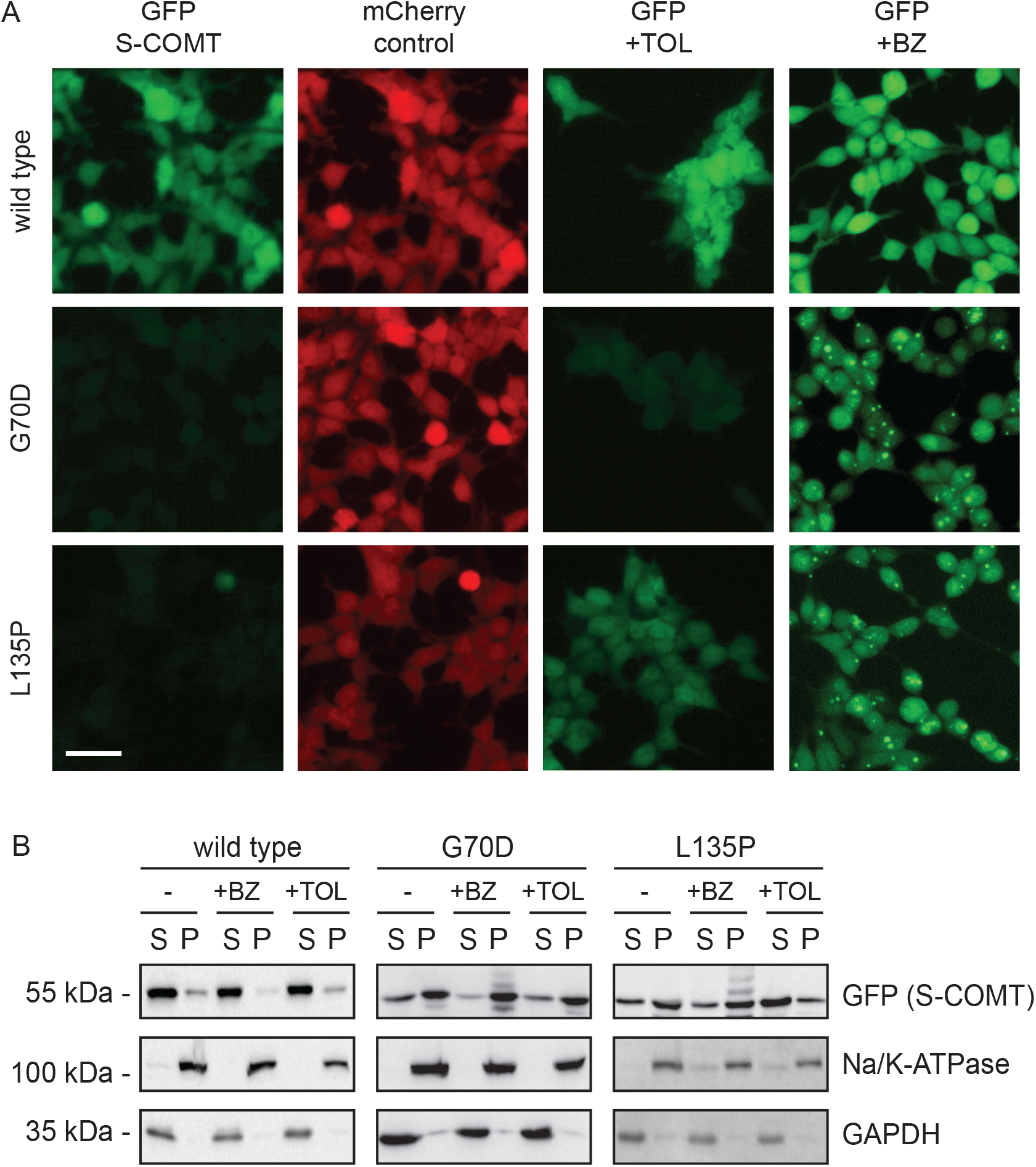
Cellular localisation and aggregation of S-COMT variants. (A) The cellular localisation of wild type (WT) S-COMT, G70D and L135P in HEK293T cells analysed by live cell fluorescence imaging. The scale bar measures 10 μm. (B) Whole cell lysates of cells expressing S-COMT WT, G70D and L135P were divided into soluble supernatant (S) or insoluble pellet (P) fractions by centrifugation after no treatment or incubation with the proteasomal inhibitor bortezomib (BZ) or the COMT inhibitor tolcapone (TOL). The protein levels in the fractions were compared by SDS-PAGE and western blotting using antibodies against GFP. Antibodies against Na/K-ATPase and GAPDH were used as controls for the insoluble and soluble fractions, respectively.

### COMT variants display reduced steady-state levels in the membrane-bound isoform

To assess the effect of COMT variants in the membrane-bound isoform, the same missense variants were generated in MB-COMT C-terminally linked to a GFP-tag and expressed from the landing-pad as above. The two isoforms have identical protein sequences, with the exception of a 50-residue N-terminal extension containing a transmembrane domain in MB-COMT. Noticeably, the reduction in steady-state level is independent of whether the variants were expressed as the membrane-bound or the soluble isoform of the protein. That is, the two MB-COMT variants, G120D and L185P (S-COMT: G70D and L135P), displayed reduced cellular abundance and MB-COMT V158M (S-COMT: V108M) displayed wild type-like abundance (Fig. 5A). These results were corroborated by flow cytometry and fluorescence microscopy (Fig. 5BC), which showed that cells expressing MB-COMT G120D and L185P displayed a tendency for reduced GFP signal and V158M displayed wild type-like GFP signal.

**Fig. 5.**
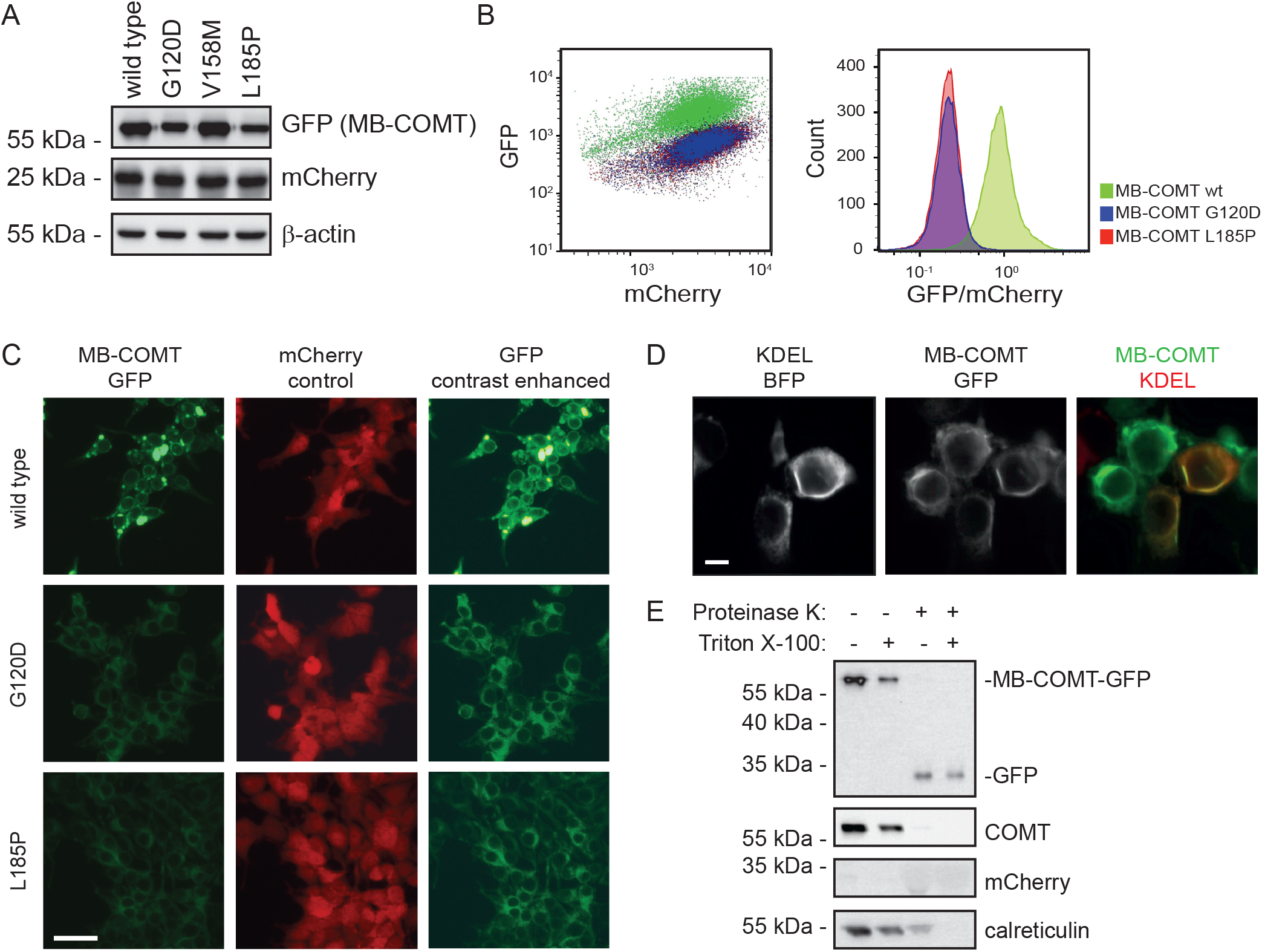
Two COMT variants are also unstable in the membrane-bound isoform. (A) The steadystate levels of whole cell lysates of HEK293T cells expressing wild type (WT) MB-COMT and the variants G120D (S-COMT: G70D) and L185P (S-COMT: L135P) determined by SDS-PAGE and western blotting using antibodies against GFP. Probing for mCherry and β-actin was included as controls for equal expression and loading, respectively. (B) Quantification by flow cytometry of the GFP and mCherry signals in HEK293T cells expressing WT MB-COMT and the two unstable variants, G120D and L185P, shown as a scatter plot (left panel) and as a histogram of the GFP/mCherry ratio (right panel). (C) The cellular localisation of WT MB-COMT, G120D and L185P in HEK293T cells analysed by live cell fluorescence imaging. The scale bar measures 10 μm. (D) Co-localisation of WT MB-COMT with the blue fluorescent protein (BFP) fused to an ER-retention sequence KDEL analysed by live cell fluorescence imaging. (E) Membrane topology of WT MB-COMT was determined by purification of microsomes from HEK293T cells expressing WT MB-COMT with a C-terminal GFP-tag following treatment with proteinase K and/or Triton X-100 as indicated. Levels of MB-COMT were determined by SDS-PAGE and western blotting using antibodies against GFP and COMT. Probing for mCherry and calreticulin was included as controls for equal expression and loading, respectively.

Since there has been some debate on the subcellular localisation of MB-COMT (Chen et al., 2011; Lundström et al., 1995; Schott et al., 2010; Ulmanen et al., 1997) we further explored the localisation of this isoform. Co-transfection with a plasmid encoding blue fluorescence protein (BFP) fused to the ER-retention sequence KDEL revealed that MB-COMT localises to the ER and thus, must be anchored in the ER membrane (Fig 5D). To investigate the topology of MB-COMT in the ER membrane, microsomes were purified and treated with proteinase K. While the ER luminal protein calreticulin showed some resistance to the treatment, the MB-COMT signal was almost entirely lost (Fig. 5E). This indicates that the GFP-tagged C-terminal of MB-COMT is oriented towards the cytosol. In addition, we note a fragment corresponding to GFP appeared resistant to proteinase K.

### Evolutionary conservation of COMT

Finally, to further assess the mutational tolerance of COMT variants, we analysed the evolutionary conservation of COMT orthologues across a broad range of species. First, we generated a multiple sequence alignment of 1490 different COMT orthologues. This was then used as an input to the GEMME model (Laine et al., 2019) that considers both residue and pair conservation to determine evolutionary distance scores (ΔE), which on the residue level reports on the likelihood of a given substitution. In this implementation, neutral variations with no or limited effects on the protein stability and/or function display ΔE scores close to zero, while substitutions with a large negative score (ΔE < −3) are likely unfavourable. As for the structural stability predictions (Fig. 1A), the ΔE scores are presented as a heat-map (Fig. 6A), and all the scores are listed in the supplemental material (Supplemental File 1). Since these evolutionary conservation scores are independent of the COMT crystal structure, the entire MB-COMT (and S-COMT) sequence is covered.

**Fig. 6.**
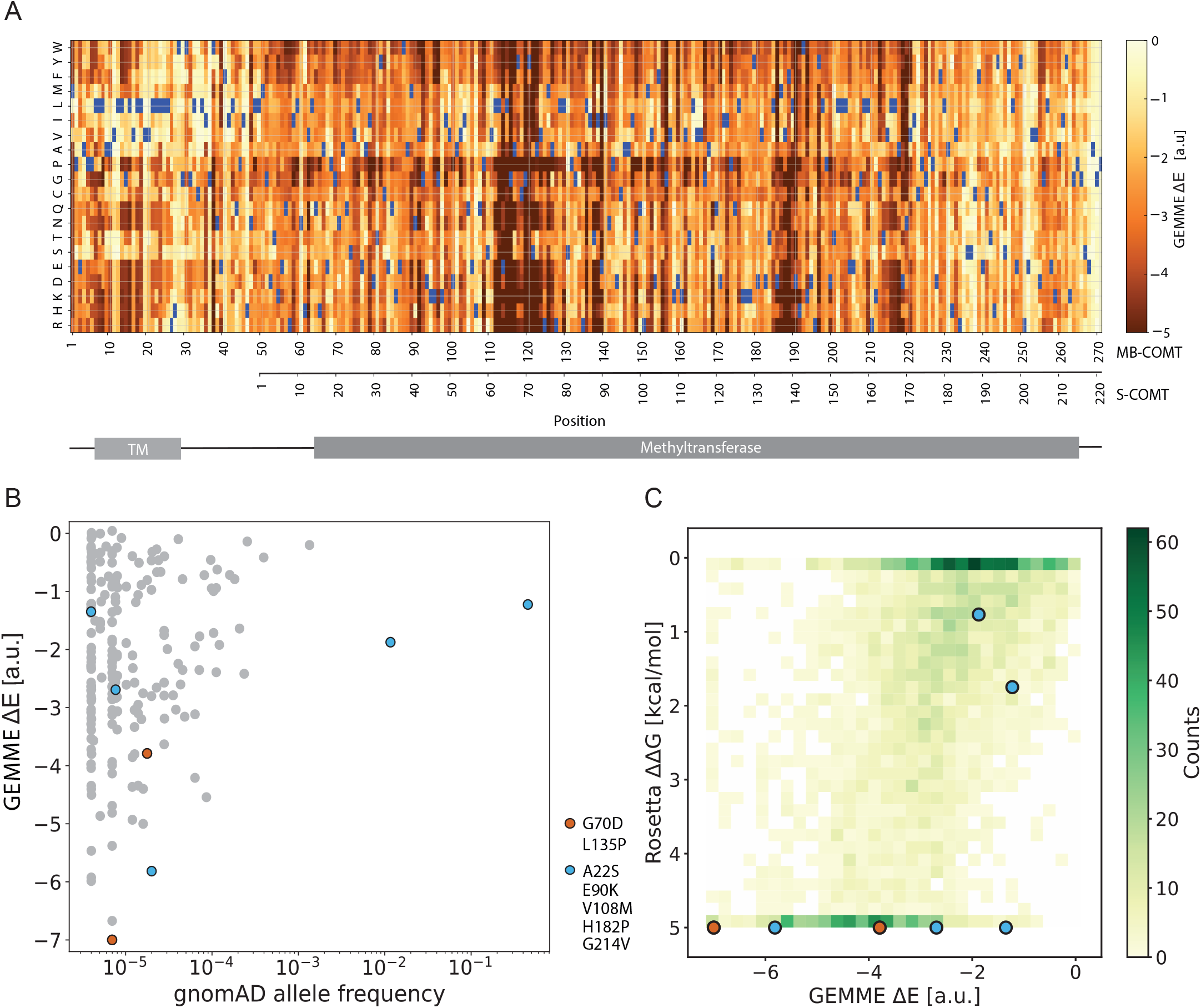
Evolutionary conservation of COMT variants. A heatmap of the evolutionary distance scores of COMT variants calculated by the GEMME model based on a multiple sequence alignment of 1490 different COMT orthologues. Substitutions with a ΔE < −3 are predicted to be detrimental. The numeration of S-COMT and MB-COMT is indicated, as well as the transmembrane (TM) and methyltransferase domain in the COMT structure (Letunic et al., 2021). (B) The GEMME ΔE scores of variants reported in the Genome Aggregation Database (gnomAD) (Karczewski et al., 2020). The two unstable variants, G70D/G120D (S-COMT/MB-COMT) and L135P/L185P (S-COMT/MB-COMT), are indicated in red and stable variants (S-COMT: A22S, E90K, V108M, H182P, G214V) in blue. (C) Mapping COMT variants by their GEMME ΔE scores and Rosetta ΔΔG scores. The two unstable variants, G70D/G120D (S-COMT/MB-COMT) and L135P/L185P (S-COMT/MB-COMT), are indicated in red and stable variants indicated in blue.

When we compared the ΔE scores with the COMT variants reported in gnomAD we observed that common alleles displayed wild type-like scores (ΔE scores close to zero) (Fig. 6BC), suggesting that they are folded and functional proteins. Conversely, the unstable G70D (G120D) and L135P (L185P), respectively, scored −7.0 and −3.8, indicating that these substitutions are detrimental.

## Discussion

It is well established that changes in our DNA can have substantial implications on our health and disease. Loss-of-function variants may decrease protein stability and/or block an interaction site or directly inhibit enzyme activity. It has been established that decreased structural stability of a protein induced by certain missense mutations may lead to rapid degradation of the protein (Stein et al., 2019). Such missense mutations have been shown to play a key role in the development of multiple diseases such as Lynch syndrome (Kampmeyer et al., 2017b; Nielsen et al., 2017), phenylketonuria (Pey et al., 2003), cystic fibrosis (Ahner et al., 2007; Meacham et al., 2001), Birt-Hogg-Dubé syndrome (Clausen et al., 2020), Parkinson’s disease (Olzmann et al., 2004) and also of importance for drug metabolism (Amorosi et al., 2021; Geck et al., 2022; Matreyek et al., 2018; Suiter et al., 2020). This has prompted a drive to understand the effects of genetic mutations on protein stability and the detailed mechanisms of the protein quality control system.

In this study, we have characterised a series of common and rare missense variants of S-COMT, focusing further on the two rare variants G70D and L135P. These two variants display severely decreased cellular steady-state levels compared to the wild type (Fig. 2CD). Treatment with the proteasome inhibitor bortezomib effectively blocked this reduction revealing that the variants are substrates of the proteasome (Fig. 2F). Additionally, we demonstrate that the variants are mainly insoluble (Fig 4B) and poly-ubiquitylated (Fig. 2H). These results correlate with the variants’ predicted ΔΔG values of 8.6 and 6.8 kcal/mol, respectively (Table 1), suggesting the variants to be structurally unstable. Upon treatment with the well-characterised inhibitors, entacapone and tolcapone (Lees, 2008), G70D is unaffected, whereas, L135P becomes stabilised and is no-longer a target of the UPS leading to an increase in the cellular levels of the variant (Fig. 3). This indicates that while the L135P variant retains its ability to bind COMT inhibitors, G70D is either unable to bind, or that the thermodynamic driving force from binding is simply insufficient to restabilize COMT. Indeed, Gly70 is located next to the SAM co-factor, which in turn interacts with the COMT inhibitors, and we suggest that G70D might affect SAM binding and thereby the ability of the COMT inhibitors to bind. Our observations are thus overall in line with previous studies suggesting that PQC-linked degradation can appear over-meticulous and may target variants that are still partly functional (Gardner et al., 2005; Kampmeyer et al., 2022; Kampmeyer et al., 2017a; Kriegenburg et al., 2014; Meacham et al., 2001).

Single-site substitutions to proline such as L135P are often detrimental, but the structural effect of such a mutation is challenging to model and predict (Alford et al., 2017; Frenz et al., 2020). However, due to its position in the beginning of a β-strand in the middle of the seven β-sheet core (Fig. S2), it is likely that the substituted proline is unable to establish the necessary hydrogen-bonds in this position causing a destabilising effect. Glycine 70 sits at the beginning of an α-helix (Fig. S2), close to the SAM binding site, and populates the so-called α_L_ -region of the Ramachandran map (with a positive φ dihedral angle). While aspartic acid may also populate the α_L_-region, it does so less favorably than glycine (Hovmöller et al., 2002). Thus, G70D might destabilize COMT via several effects including perturbing the tight turn in this position, introducing a negative charge in a partially buried region and perturbing binding of SAM.

In addition to G70D and L135P, the cellular steady-state level of five other variants was investigated, including the variants E90K and H182P. The Rosetta structural stability calculations predicted ΔΔGs of these two variants as 5.4 and 6.7 kcal/mol, respectively (Table 1), suggesting that these variants are unstable. However, both variants displayed wild type-like abundance, indicating that these variants are not targeted and subjected to UPS-mediated degradation. In the case of H182P, we speculate that the predicted destabilisation of this variant might be due to decreased accuracy when predicting changes in stability for substitutions involving proline (Frenz et al., 2020). For the E90K variant we are currently unable to explain why the Rosetta prediction fails for this substitution.

Studies have presented contradictory results regarding the cellular localisation and topology of MB-COMT. The membrane-bound isoform has been detected in various cellular compartments ranging from the cell membrane with an extracellular C-terminal orientation (Chen et al., 2011) to the membrane of the rough ER (Lundström et al., 1995; Ulmanen et al., 1997). Here we demonstrate that MB-COMT is likely anchored in the ER membrane with the C-terminal enzymatic domain oriented towards the cytosol (Fig. 5DE). Additionally, the results show that the MB-COMT variants, G120D (S-COMT: G70D) and L185P (S-COMT: L135P), are degraded. This indicates that the structural destabilisation leading to proteasomal degradation is independent of the isoform, and that any potential global structural stabilisation offered by the transmembrane domain is not sufficient to spare these variants from degradation.

The S-COMT variants, A22S and V108M, have previously been suggested to locate to a hotspot for inactivation of COMT (Rutherford & Daggett, 2009). Additionally, the level of both enzymatically active and immunoreactive V108M protein is reduced when this variant is expressed in either HEK293T and COS-1 cells (Shield et al., 2004). The intrinsic enzymatic activity of V108M is comparable to that of wild type COMT, and A22S shows a 20–30% reduced catalytic efficiency (Li et al., 2005). This suggests that missense mutations displaying reduced enzymatic activity are a result of reduced intracellular levels of the enzyme, and both variants were shown to be more thermolabile than the wild type protein (Li et al., 2005). However, in our study, both variants, A22S and V108M, were predicted to have relatively low ΔΔG values of 0.8 and 1.8 kcal/mol, respectively (Table 1), and indeed, both variants display wild type-like abundance (Fig. 2CD). This is in line with measurements of thermodynamic stability that shows that ΔΔG < 0.5 kcal/mol for V108M (Rutherford et al., 2008). Hence, these data indicate that reduced stability and degradation is not the main cause of their enzymatic inactivation.

Characterisation of COMT variants either displaying reduced enzymatic activity or rapid proteasomal degradation could potentially be of importance in Parkinson’s disease. Patients expressing these variants may be able to better compensate for the loss of dopaminergic neurons, which could possibly impact the age of disease onset. Additionally, the current symptomatic treatment of these patients includes administering the dopamine precursor, L-DOPA, together with both COMT and DDC inhibitors as adjunct drugs. However, as most COMT inhibitors are unable to cross the BBB they only provide a peripheral benefit (Müller, 2015). Therefore, patients expressing low-abundance variants or variants with reduced activity may respond better to L-DOPA treatment in that a larger amount of L-DOPA will reach the brain and the CNS due to decreased turn-over of the drug. This may allow for a reduction in the required L-DOPA administered. As treatment with L-DOPA causes multiple side-effects such as electrocardiographic changes, nausea and dyskinesia, and long-term treatment is associated with reduced efficiency of the drug (Müller, 2015), this may provide both health and economic advantages for the patient. To this end, the results presented here could provide a first step towards a personalised medicine approach with L-DOPA and COMT inhibitors.

## Supplemental information

Supplemental figures accompany this paper. The Rosetta and GEMME predictions for all COMT variants are included as a supplemental spreadsheet (Supplemental File 1).

## Acknowledgements

The authors thank Elin Josefina Pietras, Anne-Marie Lauridsen and Søren Lindemose for excellent technical assistance. We thank Anders Haagen Beck Frederiksen for assistance in early parts of the project.

## Author contributions

F.B.L., M.C., and J.D. conducted the experiments. Data analyses by F.B.L., M.C., A.S., K.L.L., and R.H.P. Experimental design by F.B.L., M.C., K.L.L., and R.H.P. K.L.L., and R.H.P. conceived the study. F.B.L. and R.H.P. wrote the paper.

## References

Abildgaard, A. B., Stein, A., Nielsen, S. V., Schultz-Knudsen, K., Papaleo, E., Shrikhande, A., Hoffmann, E. R., Bernstein, I., Anne-Marie, G., Takahashi, M., Ishioka, C., Lindorff-Larsen, K., & Hartmann-Petersen, R. (2019). Computational and cellular studies reveal structural destabilization and degradation of mlh1 variants in lynch syndrome. ELife, 8. https://doi.org/10.7554/eLife.49138

Ahner, A., Nakatsukasa, K., Zhang, H., Frizzell, R. A., & Brodsky, J. L. (2007). Small Heat-Shock Proteins Select F508-CFTR for Endoplasmic Reticulum-associated Degradation. Molecular Biology of the Cell, 18(March), 806–814. https://doi.org/10.1091/mbc.E06

Alford, R. F., Leaver-Fay, A., Jeliazkov, J. R., O’Meara, M. J., DiMaio, F. P., Park, H., Shapovalov, M. V., Renfrew, P. D., Mulligan, V. K., Kappel, K., Labonte, J. W., Pacella, M. S., Bonneau, R., Bradley, P., Dunbrack, R. L., Das, R., Baker, D., Kuhlman, B., Kortemme, T., & Gray, J. J. (2017). The Rosetta All-Atom Energy Function for Macromolecular Modeling and Design. Journal of Chemical Theory and Computation, 13(6), 3031–3048. https://doi.org/10.1021/acs.jctc.7b00125

Amorosi, C. J., Chiasson, M. A., McDonald, M. G., Wong, L. H., Sitko, K. A., Boyle, G., Kowalski, J. P., Rettie, A. E., Fowler, D. M., & Dunham, M. J. (2021). Massively parallel characterization of CYP2C9 variant enzyme activity and abundance. American Journal of Human Genetics, 108(9), 1735–1751. https://doi.org/10.1016/j.ajhg.2021.07.001

Arlow, T., Scott, K., Wagenseller, A., & Gammie, A. (2013). Proteasome inhibition rescues clinically significant unstable variants of the mismatch repair protein Msh2. Proceedings of the National Academy of Sciences of the United States of America, 110(1), 246–251. https://doi.org/10.1073/pnas.1215510110

Axelrod, J., Senoh, S., & Witkop, B. (1958). O-Methylation of catechol amines in vivo. The Journal of Biological Chemistry, 233(3), 697–701. https://doi.org/10.1016/s0021-9258(18)64730-1

Blaabjerg, L. M., Kassem, M. M., Good, L. L., Jonsson, N., Cagiada, M., Johansson, K. E., Boomsma, W., Stein, A., & Lindorff-Larsen, K. (2022). Rapid protein stability prediction using deep learning representations. BioRxiv, 2022.07.14.500157. https://www.biorxiv.org/content/10.1101/2022.07.14.500157v2%0Ahttps://www.biorxiv.org/content/10.1101/2022.07.14.500157v2.abstract

Cagiada, M., Johansson, K. E., Valanciute, A., Nielsen, S. V., Hartmann-Petersen, R., Yang, J. J., Fowler, D. M., Stein, A., & Lindorff-Larsen, K. (2021). Understanding the Origins of Loss of Protein Function by Analyzing the Effects of Thousands of Variants on Activity and Abundance. Molecular Biology and Evolution, 38(8), 3235–3246. https://doi.org/10.1093/molbev/msab095

Chen, J., Song, J., Yuan, P., Tian, Q., Ji, Y., Ren-Patterson, R., Liu, G., Sei, Y., & Weinberger, D. R. (2011). Orientation and cellular distribution of membrane-bound catechol-O-methyltransferase in cortical neurons: Implications for drug development. Journal of Biological Chemistry, 286(40), 34752–34760. https://doi.org/10.1074/jbc.M111.262790

Chi Htun, N., Miyaki, K., Song, Y., Ikeda, S., Shimbo, T., & Muramatsu, M. (2011). Association of the catechol-O-methyl transferase gene Val158Met polymorphism with blood pressure and prevalence of hypertension: Interaction with dietary energy intake. American Journal of Hypertension, 24(9), 1022–1026. https://doi.org/10.1038/ajh.2011.93

Clausen, L., Stein, A., Grønbæk-Thygesen, M., Nygaard, L., Søltoft, C. L., Nielsen, S. V., Lisby, M., Ravid, T., Lindorff-Larsen, K., & Hartmann-Petersen, R. (2020). Folliculin variants linked to Birt-Hogg-Dubésyndrome are targeted for proteasomal degradation. PLoS Genetics, 16(11). https://doi.org/10.1371/journal.pgen.1009187

Dextera, D. T., & Jenner, P. (2013). Parkinson disease: From pathology to molecular disease mechanisms. Free Radical Biology and Medicine, 62, 132–144. https://doi.org/10.1016/j.freeradbiomed.2013.01.018

Ehler, A., Benz, J., Schlatter, D., & Rudolph, M. G. (2014). Mapping the conformational space accessible to catechol-O-methyltransferase. Acta Crystallographica Section D: Biological Crystallography, 70(8), 2163–2174. https://doi.org/10.1107/S1399004714012917

Frank, M. J., & Fossella, J. A. (2011). Neurogenetics and pharmacology of learning, motivation, and cognition. Neuropsychopharmacology, 36(1), 133–152. https://doi.org/10.1038/npp.2010.96

Frenz, B., Lewis, S. M., King, I., DiMaio, F., Park, H., & Song, Y. (2020). Prediction of Protein Mutational Free Energy: Benchmark and Sampling Improvements Increase Classification Accuracy. Frontiers in Bioengineering and Biotechnology, 8(October), 1–8. https://doi.org/10.3389/fbioe.2020.558247

Friedman, J. R., Lackner, L. L., West, M., DiBenedetto, J. R., Nunnari, J., & Voeltz, G. K. (2011). ER tubules mark sites of mitochondrial division. Science, 334(6054), 358–362. https://doi.org/10.1126/science.1207385

Gardner, R. G., Nelson, Z. W., & Gottschling, D. E. (2005). Degradation-mediated protein quality control in the nucleus. Cell, 120(6), 803–815. https://doi.org/10.1016/j.cell.2005.01.016

Gatt, J. M., Burton, K. L. O., Williams, L. M., & Schofield, P. R. (2015). Specific and common genes implicated across major mental disorders: A review of meta-analysis studies. Journal of Psychiatric Research, 60, 1–13. https://doi.org/10.1016/j.jpsychires.2014.09.014

Geck, R. C., Boyle, G., Amorosi, C. J., Fowler, D. M., & Dunham, M. J. (2022). Measuring Pharmacogene Variant Function at Scale Using Multiplexed Assays. Annual Review of Pharmacology and Toxicology, 62, 531–550. https://doi.org/10.1146/annurev-pharmtox-032221-085807

Gray, V. E., Hause, R. J., & Fowler, D. M. (2017). Analysis of large-scale mutagenesis data to assess the impact of single amino acid substitutions. Genetics, 207(1), 53–61. https://doi.org/10.1534/genetics.117.300064

Hanna, C. (1965). Metabolism of catecholamines. Investigative Ophthalmology, 4(6), 1095–1104. https://doi.org/10.1007/978-1-4615-7163-6_9

Høie, M. H., Cagiada, M., Beck Frederiksen, A. H., Stein, A., & Lindorff-Larsen, K. (2022). Predicting and interpreting large-scale mutagenesis data using analyses of protein stability and conservation. Cell Reports, 38(2), 110207. https://doi.org/10.1016/j.celrep.2021.110207

Hovmöller, S., Zhou, T., & Ohlson, T. (2002). Conformations of amino acids in proteins. Acta Crystallographica Section D: Biological Crystallography, 58(5), 768–776. https://doi.org/10.1107/S0907444902003359

Kampmeyer, C., Karakostova, A., Schenstrøm, S. M., Abildgaard, A. B., Lauridsen, A. M., Jourdain, I., & Hartmann-Petersen, R. (2017). The exocyst subunit Sec3 is regulated by a protein quality control pathway. Journal of Biological Chemistry, 292(37), 15240–15253. https://doi.org/10.1074/jbc.M117.789867

Kampmeyer, C., Larsen-Ledet, S., Wagnkilde, M. R., Michelsen, M., Iversen, H. K. M., Nielsen, S. V., Lindemose, S., Caregnato, A., Ravid, T., Stein, A., Teilum, K., Lindorff-Larsen, K., & Hartmann-Petersen, R. (2022). Disease-linked mutations cause exposure of a protein quality control degron. Structure, 30(9), 1245–1253.e5. https://doi.org/10.1016/j.str.2022.05.016

Kampmeyer, C., Nielsen, S. V., Clausen, L., Stein, A., Gerdes, A. M., Lindorff-Larsen, K., & Hartmann-Petersen, R. (2017). Blocking protein quality control to counter hereditary cancers. Genes Chromosomes and Cancer, 56(12), 823–831. https://doi.org/10.1002/gcc.22487

Karczewski, K. J., Francioli, L. C., Tiao, G., Cummings, B. B., Alföldi, J., Wang, Q., Collins, R. L., Laricchia, K. M., Ganna, A., Birnbaum, D. P., Gauthier, L. D., Brand, H., Solomonson, M., Watts, N. A., Rhodes, D., Singer-Berk, M., England, E. M., Seaby, E. G., Kosmicki, J. A., … MacArthur, D. G. (2020). The mutational constraint spectrum quantified from variation in 141,456 humans. Nature, 581(7809), 434–443. https://doi.org/10.1038/s41586-020-2308-7

Krauß, J., & Bracher, F. (2018). Pharmacokinetic enhancers (Boosters)—escort for drugs against degrading enzymes and beyond. Scientia Pharmaceutica, 86(4). https://doi.org/10.3390/scipharm86040043

Kriegenburg, F., Jakopec, V., Poulsen, E. G., Nielsen, S. V., Roguev, A., Krogan, N., Gordon, C., Fleig, U., & Hartmann-Petersen, R. (2014). A Chaperone-Assisted Degradation Pathway Targets Kinetochore Proteins to Ensure Genome Stability. PLoS Genetics, 10(1). https://doi.org/10.1371/journal.pgen.1004140

Lachman, H. M., Papolos, D. F., Saito, T., Yu, Y. M., Szumlanski, C. L., & Weinshilboum, R. M. (1996). Human catechol-O-methyltransferase pharmacogenetics: Description of a functional polymorphism and its potential application to neuropsychiatric disorders [Article]. Pharmacogenetics, 6(3), 243–250. https://doi.org/10.1097/00008571-199606000-00007

Laine, E., Karami, Y., & Carbone, A. (2019). GEMME: A Simple and Fast Global Epistatic Model Predicting Mutational Effects. Molecular Biology and Evolution, 36(11), 2604–2619. https://doi.org/10.1093/molbev/msz179

Lees, A. J. (2008). Evidence-Based Efficacy Comparison of Tolcapone and Entacapone as Adjunctive Therapy in Parkinson’s Disease. CNS Drug Reviews, 14(1), 83–93. https://doi.org/10.1111/j.1527-3458.2007.00035.x

Letunic, I., Khedkar, S., & Bork, P. (2021). SMART: Recent updates, new developments and status in 2020. Nucleic Acids Research, 49(D1), D458–D460. https://doi.org/10.1093/nar/gkaa937

Li, Y., Yang, X., Van Breemen, R. B., & Bolton, J. L. (2005). Characterization of two new variants of human catechol O-methyltransferase in vitro. Cancer Letters, 230(1), 81–89. https://doi.org/10.1016/j.canlet.2004.12.022

Lundström, K., Tenhunen, J., Tilgmann, C., Karhunen, T., Panula, P., & Ulmanen, I. (1995). Cloning, expression and structure of catechol-O-methyltransferase. J. Biochem. Biophys., 1251, 1–10.

Männistö, P. T., & Kaakkola, S. (1999). Catechol-O-methyltransferase (COMT): Biochemistry, molecular biology, pharmacology, and clinical efficacy of the new selective COMT inhibitors. Pharmacological Reviews, 51(4), 593–628.

Matreyek, K. A., Starita, L. M., Stephany, J. J., Martin, B., Chiasson, M. A., Gray, V. E., Kircher, M., Khechaduri, A., Dines, J. N., Hause, R. J., Bhatia, S., Evans, W. E., Relling, M. V., Yang, W., Shendure, J., & Fowler, D. M. (2018). Multiplex assessment of protein variant abundance by massively parallel sequencing. Nature Genetics, 50(6), 874–882. https://doi.org/10.1038/s41588-018-0122-z

Matreyek, K. A., Stephany, J. J., Chiasson, M. A., Hasle, N., & Fowler, D. M. (2020). An improved platform for functional assessment of large protein libraries in mammalian cells. Nucleic Acids Research, 48(1), 1–12. https://doi.org/10.1093/nar/gkz910

Meacham, G. C., Patterson, C., Zhang, W., Younger, J. M., & Cyr, D. M. (2001). The Hsc70 co-chaperone CHIP targets immature CFTR for proteasomal degradation. Nature Cell Biology, 3(1), 100–105. https://doi.org/10.1038/35050509

Moo-On Huh, M., & Friedhoff, A. J. (1979). Multiple molecular forms of catechol-O-methyltransferase. Evidence for two distinct forms, and their purification and physical characterization. Journal of Biological Chemistry, 254(2), 299–308. https://doi.org/10.1016/s0021-9258(17)37918-8

Müller, T. (2015). Catechol-O-methyltransferase inhibitors in Parkinson’s disease. Drugs, 75(2), 157–174. https://doi.org/10.1007/s40265-014-0343-0

Nielsen, S. V., Hartmann-Petersen, R., Stein, A., & Lindorff-Larsen, K. (2021). Multiplexed assays reveal effects of missense variants in MSH2 and cancer predisposition. PLoS Genetics, 17(4 April 2021). https://doi.org/10.1371/journal.pgen.1009496

Nielsen, S. V., Poulsen, E. G., Rebula, C. A., & Hartmann-Petersen, R. (2014). Protein quality control in the nucleus. Biomolecules, 4(3), 646–661. https://doi.org/10.3390/biom4030646

Nielsen, S. V., Stein, A., Dinitzen, A. B., Papaleo, E., Tatham, M. H., Poulsen, E. G., Kassem, M. M., Rasmussen, L. J., Lindorff-Larsen, K., & Hartmann-Petersen, R. (2017). Predicting the impact of Lynch syndrome-causing missense mutations from structural calculations. PLOS Genetics, 13(4), 1–26. https://doi.org/10.1371/journal.pgen.1006739

Olzmann, J. A., Brown, K., Wilkinson, K. D., Rees, H. D., Huai, Q., Ke, H., Levey, A. I., Li, L., & Chin, L. S. (2004). Familial Parkinson’s Disease-associated L166P Mutation Disrupts DJ-1 Protein Folding and Function. Journal of Biological Chemistry, 279(9), 8506–8515. https://doi.org/10.1074/jbc.M311017200

Park, H., Bradley, P., Greisen, P., Liu, Y., Mulligan, V. K., Kim, D. E., Baker, D., & Dimaio, F. (2016). Simultaneous Optimization of Biomolecular Energy Functions on Features from Small Molecules and Macromolecules. Journal of Chemical Theory and Computation, 12(12), 6201–6212. https://doi.org/10.1021/acs.jctc.6b00819

Pey, A. L., Desviat, L. R., Gámez, A., Ugarte, M., & Pérez, B. (2003). Phenylketonuria: Genotypephenotype correlations based on expression analysis of structural and functional mutations in PAH. Human Mutation, 21(4), 370–378. https://doi.org/10.1002/humu.10198

Remmert, M., Biegert, A., Hauser, A., & Söding, J. (2012). HHblits: Lightning-fast iterative protein sequence searching by HMM-HMM alignment. Nature Methods, 9(2), 173–175. https://doi.org/10.1038/nmeth.1818

Rutherford, K., Alphandéry, E., McMillan, A., Daggett, V., & Parson, W. W. (2008). The V108M mutation decreases the structural stability of catechol O-methyltransferase. Biochimica et Biophysica Acta - Proteins and Proteomics, 1784(7–8), 1098–1105. https://doi.org/10.1016/j.bbapap.2008.04.006

Rutherford, K., & Daggett, V. (2009). A hotspot of inactivation: The A22S and V108M polymorphisms individually destabilize the active site structure of catechol O-methyltransferase. Biochemistry, 48(27), 6450–6460. https://doi.org/10.1021/bi900174v

Schott, B. H., Frischknecht, R., Debska-Vielhaber, G., John, N., Behnisch, G., Düzel, E., Gundelfinger, E. D., & Seidenbecher, C. I. (2010). Membrane-bound catechol-O-methyl transferase in cortical neurons and glial cells is intracellularly oriented. Frontiers in Psychiatry, 1(OCT), 1–9. https://doi.org/10.3389/fpsyt.2010.00142

Shield, A. J., Thomae, B. A., Eckloff, B. W., Wieben, E. D., & Weinshilboum, R. M. (2004). Human catechol O-methyltransferase genetic variation: Gene resequencing and functional characterization of variant allozymes. Molecular Psychiatry, 9(2), 151–160. https://doi.org/10.1038/sj.mp.4001386

Stein, A., Fowler, D. M., Hartmann-Petersen, R., & Lindorff-Larsen, K. (2019). Biophysical and Mechanistic Models for Disease-Causing Protein Variants. Trends in Biochemical Sciences, 44(7), 575–588. https://doi.org/10.1016/j.tibs.2019.01.003

Suiter, C. C., Moriyama, T., Matreyek, K. A., Yang, W., Scaletti, E. R., Nishii, R., Yang, W., Hoshitsuki, K., Singh, M., Trehan, A., Parish, C., Smith, C., Li, L., Bhojwani, D., Yuen, L. Y. P., Li, C. kong, Li, C. ho, Yang, Y. li, Walker, G. J., … Yang, J. J. (2020). Massively parallel variant characterization identifies NUDT15 alleles associated with thiopurine toxicity. Proceedings of the National Academy of Sciences of the United States of America, 117(10), 5394–5401. https://doi.org/10.1073/pnas.1915680117

Teng, Y., He, C., Zuo, X., & Li, X. (2013). Catechol-O-methyltransferase and cytochrome P-450 1B1 polymorphisms and endometrial cancer risk□: A meta-analysis. International Journal of Gynecological Cancer, 23(3), 422–430. https://doi.org/10.1097/IGC.0b013e3182849e0d

Tenhunen, J., Salminen, M., Lundström, K., Kiviluoto, T., Savolainen, R., & Ulmanen, I. (1994). Genomic organization of the human catechol O□methyltransferase gene and its expression from two distinct promoters. European Journal of Biochemistry, 223(3), 1049–1059. https://doi.org/10.1111/j.1432-1033.1994.tb19083.x

Ulmanen, I., Peränen, J., Tenhunen, J., Tilgmann, C., Karhunen, T., Panula, P., Bernasconi, L., Aubry, J. P., & Lundström, K. (1997). Expression and intracellular localization of catechol O-methyltransferase in transfected mammalian cells. European Journal of Biochemistry, 243(1–2), 452–459. https://doi.org/10.1111/j.1432-1033.1997.0452a.x

Voutilainen, S., Tuomainen, T. P., Korhonen, M., Mursu, J., Virtanen, J. K., Happonen, P., Alfthan, G., Erlund, I., North, K. E., Mosher, M. J., Kauhanen, J., Tiihonen, J., Kaplan, G. A., & Salonen, J. T. (2007). Functional COMT Val158Met polymorphism, risk of acute coronary events and serum homocysteine: The Kuopio Ischaemic Heart Disease Risk Factor Study. PLoS ONE, 2(1), 1–7. https://doi.org/10.1371/journal.pone.0000181

Xiao, L., Tong, M., Jin, Y., Huang, W., & Li, Z. (2013). The l58Val/Met polymorphism of catechol-O-methyl transferase gene and prostate cancer risk: A meta-analysis. Molecular Biology Reports, 40(2), 1835–1841. https://doi.org/10.1007/s11033-012-2238-z

Yilmaz, A., & Çetin, İ. (2020). In Silico Prediction of the Effects of Nonsynonymous Single Nucleotide Polymorphisms in the Human Catechol-O-Methyltransferase (COMT) Gene. Cell Biochemistry and Biophysics, 78(2), 227–239. https://doi.org/10.1007/s12013-020-00905-6

