## Supplemental Figures for "Rare catechol-O-methyltransferase (COMT) missense variants are structurally unstable proteasome targets"

***Supplemental material***


**Figure S1.** C-terminally tagged COMT variants. p. 2

**Figure S2.** Selected variants mapped on the COMT structure. p. 3

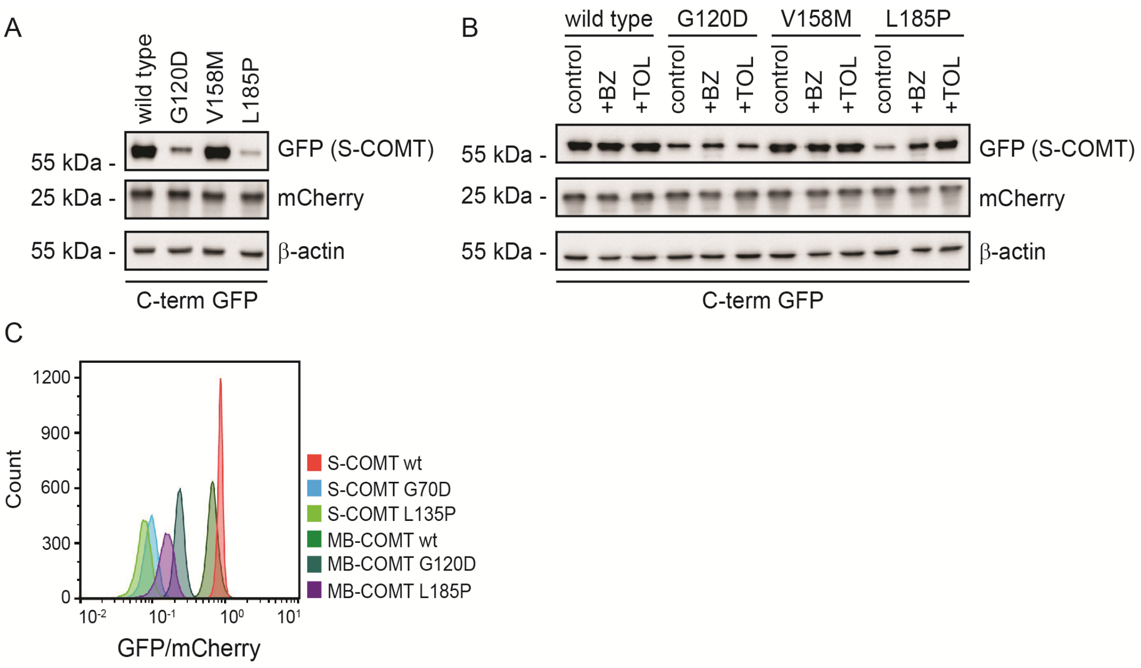


**Fig. S1 C-terminally tagged COMT variants.** (AB) The steady-state levels of whole cell lysates of HEK293T cells expressing wild type (WT) S-COMT and the variants G70D and L135P C-terminally tagged with GFP either untreated (A), treated with bortezomib (BZ) (16 hrs) or tolcapone (TOL) (24 hrs) (B) determined by SDS-PAGE and western blotting using antibodies against GFP. Antibodies against mCherry and β-actin serve as controls. (C) Quantification by flow cytometry of GFP and mCherry signal in HEK293T cells expressing WT S-COMT and the two unstable variants, G70D and L135P, C-terminally tagged with GFP.

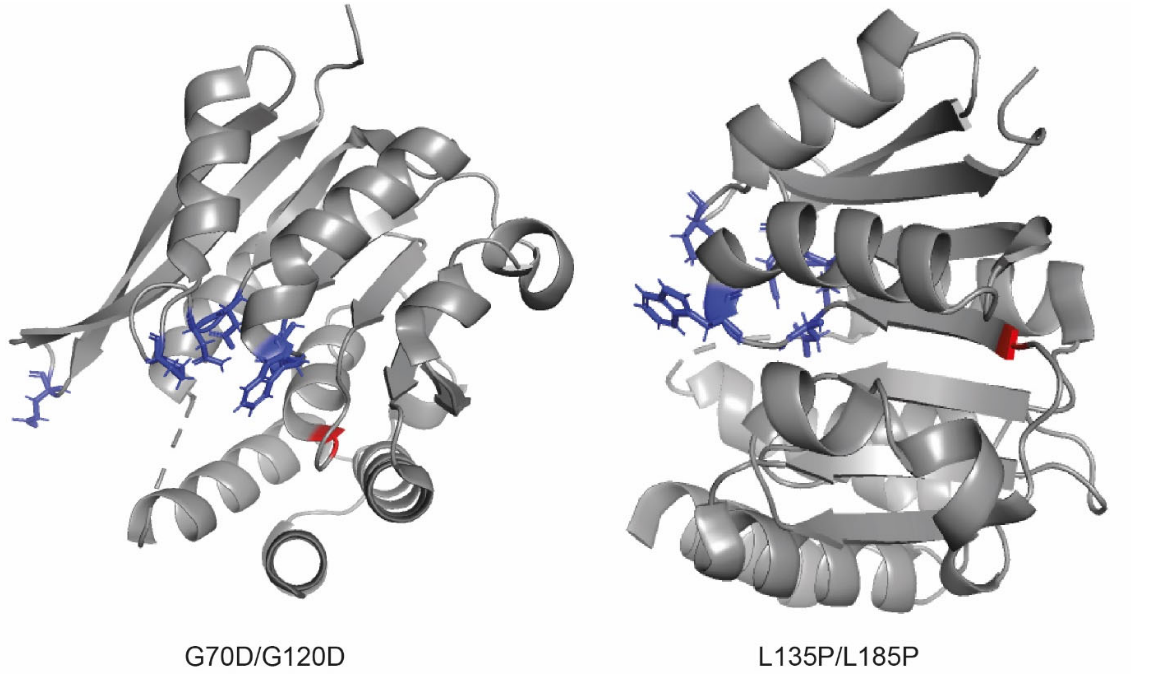


**Fig. S2 Selected variants mapped on the COMT structure.** The crystal structure of S-COMT (PDB: 4PYI) variants G70D/G120D (S-COMT/MB-COMT) (left) and L135P/L185P (S-COMT/MB-COMT) (right). Mutated residues are marked with red. Residues contributing to substrate binding and the active site are marked with blue (W88, D191, W193, D219, N220, P224, E249). The figures were created in PyMol.
